# A Machine Learning Approach for Real-time Cortical State Estimation

**DOI:** 10.1101/2023.06.20.545785

**Authors:** David A Weiss, Adriano MF Borsa, Aurélie Pala, Audrey J Sederberg, Garrett B Stanley

## Abstract

**Objective:** Cortical function is under constant modulation by internally-driven, latent variables that regulate excitability, collectively known as “cortical state”. Despite a vast literature in this area, the estimation of cortical state remains relatively ad hoc, and not amenable to real-time implementation. Here, we implement robust, data-driven, and fast algorithms that address several technical challenges for online cortical state estimation.

**Approach:** We use unsupervised Gaussian Mixture Models (GMMs) to identify discrete, emergent clusters in spontaneous local field potential (LFP) signals in cortex. We then extend our approach to a temporally-informed Hidden semi-Markov Model (HSMM) with Gaussian observations to better model and infer cortical state transitions. Finally, we implement our HSMM cortical state inference algorithms in a real-time system, evaluating their performance in emulation experiments.

**Main results:** Unsupervised clustering approaches reveal emergent state-like structure in spontaneous electrophysiological data that recapitulate arousal-related cortical states as indexed by behavioral indicators. HSMMs enable cortical state inferences in a real-time context by modeling the temporal dynamics of cortical state switching. Using HSMMs provides robustness to state estimates arising from noisy, sequential electrophysiological data.

**Significance:** To our knowledge, this work represents the first implementation of a real-time software tool for continuously decoding cortical states with high temporal resolution (40 ms). The software tools that we provide can facilitate our understanding of how cortical states dynamically modulate cortical function on a moment-by-moment basis and provide a basis for state-aware brain machine interfaces across health and disease.

## 1. Introduction

Even in the absence of exogeneous sensory inputs or volitional movement, the brain is spontaneously active, undergoing continual changes in gross activity that is thought to be representative of the state of the brain. There is ample evidence for discrete state switching in a variety of neurological contexts across health and disease. Notably, the switching between REM and NREM sleep states, each with their own distinct cortical signatures, has long been documented [1,2]. Importantly, the notion of distinct states extends to the awake brain as well. In some of the earliest human EEG recordings, Hans Berger found that having subjects perform mental arithmetic blocked prominent ∼10 Hz alpha rhythms, giving the first signs that there are distinct states of wakefulness detectable in cortex [3–5]. Discrete neural states, as indexed by oscillatory activity, have their place in neuropathophysiology too, with abnormal oscillatory states being associated with movement deficits in Parkinson’s disease [6–8] as well as transient cognitive deficits in absence epilepsy [9,10]. Indeed, switching between discrete states of activity is a ubiquitous motif in neurological systems with profound functional consequences. Despite this fact, approaches to decode brain state remain relatively ad hoc.

In the past few decades, there has been an increased push towards identifying the characteristics, mechanisms, and functional consequences of brain states as they pertain to wakefulness and arousal [11–15]. The mouse neocortex during wakefulness has proven to be a particularly fruitful model for such efforts, with many reports and characterizations of activated versus deactivated cortical states closely associated with accessible behavioral readouts [13,16–22]. Generally, animals that are actively exploring their environment or aroused (as indicated by whisking, locomotion, and/or increasing pupil size) exhibit prominent high-frequency (HF; 30-90 Hz) activity in their cortical local field potential (LFP), or an *activated* cortical state. Meanwhile, periods of quiet wakefulness with no behavioral markers of arousal are characterized by elevated low-frequency (LF; 1-10 Hz) oscillations, or a *deactivated* cortical state. Here, it is important to note the distinction between behavioral markers indexing cortical states and the cortical states themselves, which are defined in terms of underlying neural activity patterns. Already, significant progress has been made in identifying the circuit mechanisms [22–26] and the profound sensory consequences [16,19,27–29] of these cortical states.

However, to our knowledge it remains an open challenge to robustly decode cortical states in real-time. Previous studies have primarily focused on post hoc analysis of cortical states during continuous periods typically lasting several seconds [16,18,19], without decoding cortical states in uncertain regimes, such as during sub-second state fluctuations or transitioning periods between cortical states. While appropriate for post hoc analysis, such studies circumvent precisely timing the transitions between states as is needed for a continuous, real-time readout of cortical state. Additionally, cortical states are typically separated by ad hoc methods that are not robust to interindividual variability in the frequency content of behavioral states across sleep and wakefulness [30], which limits their use in robust, real-time processes. For example, a fixed threshold on the LF/HF ratio, meant to characterize the relative amplitude of LFP activity in a LF band relative to that in a HF band, is frequently used to parse between activated and deactivated cortical states [13,18]. There is much speculation that the number and nature of observed cortical states will differ based on an animal’s experience and behavioral context [12,13], further highlighting the need for more flexible methods of identifying cortical states. To address this gap, we propose a data-driven framework for providing a continuous readout of brain states that leverages the temporal dynamics of cortical state transitions and is feasible for online implementation. Real-time cortical state decoders are a promising tool to further elucidate how cortical states dynamically modulate cortical function and provide the basis for state-aware brain-machine interfaces in health and disease.

Here, we utilize machine learning approaches for identifying emergent, state-like structure in the spontaneous LFP of the whisker region of primary somatosensory cortex (S1) of the awake, head-fixed mouse. Simultaneously, we collect behavioral videography of whisking to serve as external validation for cortical states estimated solely on electrophysiological data. We first show that unsupervised Gaussian Mixture Models (GMMs) find emergent clusters that align with whisking versus quiescent periods. Then, we use temporally-informed Hidden semi-Markov Models (HSMMs) to better capture cortical state switching. Finally, we leverage the fact that the LFP is a real-time accessible signal to implement these state decoding algorithms in an online setting, for use in tracking cortical states under a variety of experimental contexts. Taken together, these approaches constitute a method for tracking temporally fluctuating brain states in a robust manner across neural and behavioral contexts.

## 2. Methods

All procedures were approved by the Institutional Animal Care and Use Committee at the Georgia Institute of Technology and were in agreement with guidelines established by the National Institutes of Health.

### 2.1 Animal preparation for head-restrained awake recordings

Ten male 8- to 26-week-old C57BL/6J mice were housed under a reversed light-dark cycle. Mice were implanted with a custom-made stainless steel headpost and a recording chamber under 1-2% isoflurane anesthesia. Mice were allowed to recover for at least 3 days prior to additional handling. Mice were then habituated to head fixation and paw restraint for 3-6 days before proceeding to awake, head-restrained electrophysiological recordings [31].

### 2.2 Silicon probe recordings

Prior to recordings, we functionally identified putative whisker columns in S1 via intrinsic optical signal imaging through a thinned skull under 1-1.25% isoflurane anesthesia [32,33]. On the day of the first recordings, mice were anesthetized (1-1.5% isoflurane anesthesia), and a small craniotomy was made over the center of a putative cortical column previously identified by intrinsic imaging. The craniotomy was then covered with silicone elastomer (Kwik-Cast, WPI), and mice were returned to their home cage for at least 2 hours to recover from anesthesia. Mice were placed on the recording setup, the silicone elastomer removed, and a 32-channel laminar silicon probe (A1×32-5 mm-25-177-A32, 25 µm interchannel spacing, Neuronexus) was slowly inserted through the craniotomy using a micromanipulator (Luigs & Neumann) at an insertion angle of 35° from the vertical and to a target depth of 1000-1100 µm [31]. All silicon probes were electrochemically plated with a poly(3,4-ethylenedioxythiophene) polymer [34,35] using a NanoZ device (White Matter) to achieve 1 kHz impedance values between 0.2 and 1 MΩ. Data collection started after a minimum of 30 minutes to allow for relaxation of the brain tissue around the probe. Continuous signals were acquired using a Cerebus (Blackrock Microsystems) acquisition system. Signals were amplified and filtered between 0.3 Hz and 7.5 kHz and digitized at 30 kHz. After the first recording session, we removed the probe and resealed the recording well with silicone elastomer. In a subset of mice, recordings were conducted in the same craniotomy across 2 consecutive days. During recording sessions whiskers were stimulated with a computer-controlled galvanometer (Cambridge Technologies) with an attached tube to stimulate individual whiskers [36].

### 2.3 LFP preprocessing

Electrophysiological traces were down-sampled to 2 kHz to obtain LFP signals. We extracted spontaneous epochs (i.e., epochs without whisker stimulation) from the recorded activity. Each recording contained periods of >10s without any whisker stimulation, for which the entire period >1s after the last delivered whisker stimulus in a train was considered spontaneous. For all analyses, we used the LFP from the middle of cortical layer 4 (L4), which was identified with the LFP, current source density (CSD), and multiunit activity (MUA) responses evoked by contralateral whisker stimuli as previously described [31]. Fourier amplitude spectra were extracted from LFP traces by taking the magnitude of the Fast Fourier Transform (FFT), for 1.024-s (2048 point) windows with a 40-ms step length, after applying a Hamming window of the corresponding length. For all analyses, only the physiological range of <100 Hz (excluding DC) was considered.

### 2.4 Labeling behavioral states

The primary goal of this study was to develop unsupervised approaches to labeling cortical state from electrophysiological data. However, as a benchmark for the unsupervised approaches, we used behavioral videography of whisking to establish a proxy label for cortical state. Whisking has been strongly tied to short time-scale fluctuations in cortical sensory processing states, providing a strong indicator of underlying cortical state [13,17,19,37]. Thus, in all experimental sessions we recorded whisker activity for post hoc analysis of the quality of our cortical state estimates. Videography was acquired from above the face for two non-trimmed whiskers on each side of the face at 200 Hz with a resolution of 14.4 pixels/mm (EoSens CL MC1362, Mikrotron). Using DeepLabCut [38] we tracked the position of each untrimmed whisker and calculated their velocities with custom routines written in MATLAB (MathWorks). The resulting time series was then smoothed and a user-defined binary threshold was used to assign “whisking” or “quiescent” labels to each time point. Behavioral labels were up-sampled to match the sampling rate of the electrophysiology and, similarly to the electrophysiology data, were binned using a 1.024-s (2048 point) wide majority vote window with 40 ms step lengths. Therefore, for every FFT observation of the LFP we had a corresponding behavioral label.

### 2.5 Dimensionality reduction of the Fourier spectra

#### 2.5.1 Canonical LF (1-10 Hz) and HF (30-90 Hz) bands

For each frequency band, the mean Low Frequency (LF) and High Frequency (HF) amplitudes were taken, yielding 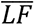 and 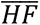 respectively [18]. We utilized a log transform on the LF/HF ratio, defined as

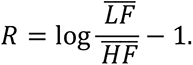

The subtraction of unity shifts the measurement so that the widely used cutoff of 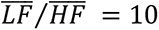 [13,18] is mapped to R = 0. When considering the frequency bands individually, the base-10 logarithm of 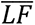 or 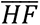 was taken as appropriate. The properties of logarithms allow thresholds on *R* to be visualized as a 45-degree line in the log HF versus log LF space.

#### 2.5.2 Principal Components Analysis (PCA)

To extend the frequency domain representation of the LFP beyond power in the classical bands, principal components analysis (PCA) was used to identify important features of the Fourier spectrum. Fourier spectra (<100 Hz, no DC) were normalized by taking the base-10 logarithm, followed by Z-scoring. PCA was then applied with a machine learning package implemented in Python (*scikit-learn*). PCA analysis yielded N feature vectors which were subsequently ℓ1-normalized, then multiplied with the normalized Fourier spectra to produce N-dimensional representations of the data. PCA features and normalization parameters were fit on training sets and used to produce PC observations from testing set data.

The features found by PCA were compared to those arising from neighborhood components analysis (NCA), which is a supervised dimensionality reduction method that finds a linear mapping to N directions which maximizes the accuracy of a stochastic K-nearest neighbors classifier [39].

### 2.6 Inferring cortical states

The dataset was split for model training and validation. Because some of our models leverage temporal information for state estimation, the first 50% of each recording was used as a training set and the last 50% for the testing set. The length of the extracted spontaneous epochs ranged from 6.5 min to 26 min per recording. Each dataset contained time evolved Fourier spectra (<100 Hz, excluding DC) extracted from the LFP of the most central L4 channel as well as cortical state labels extracted from videography. All feature identification, hyperparameter tuning, and training were conducted on the training set, with the testing set held out until final validation.

#### 2.6.1 Observations

In our experiments we utilized 1D and 2D observation models. All observations for our models were derived from frequency content of the LFP signal. We fit 1D models on the log transformed LF/HF ratio (R) and 2D models to the joint distribution of first and second principal components of the Fourier amplitude spectra (<100 Hz, excluding DC).

#### 2.6.2 One-dimensional models

First, we applied a fixed threshold on *R* to classify cortical states. Previous studies have suggested a threshold of LF/HF = 10 [13], which corresponds to a threshold in the transformed ratio of *R* = 0. Therefore, *R* < 0 corresponds to low LF activity and/or high HF activity, and an activated cortical state. Conversely, *R* > 0 corresponds to high LF activity and/or low HF activity, and a deactivated cortical state.

We also utilized a finite state Hidden Markov Model (HMM) [40,41] with univariate Gaussian observations of R to estimate underlying cortical states. In this finite state space model, the state is defined as *x* ∈ {*z_1_*, …, *z_k_*}, where *K* is the number of states. When *x* = *z_k_*, then 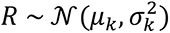. The full definition of the Hidden Markov Model is given by 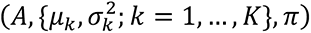, where A is a *K* × *K* matrix of state transition probabilities, *μ_k_* and 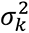 are the mean and variance of *R* while the system is in state *z_k_*, and π is the initial state distribution [41]. These parameters were fit in a maximum likelihood sense through iterative re-estimation using the Baum-Welch algorithm [42] on the training set of each recording. Once the model is learned, the most likely state sequence underlying the sequence of observations can be decoded using the Viterbi algorithm [43].

#### 2.6.3 Two-dimensional models

2D models were fit on joint distributions of the first and second PC scores with a multivariate Gaussian observation model, initially with 2D Gaussian Mixture Models (GMMs) and subsequently with Hidden Markov Models (HMMs). As in the univariate case, when the system is in state *x* = *z_k_*, then *y* ∼ *N*(*µ_k_*, *Σ_k_*), where y is now a 2D observation vector, *µ_k_* and Σ_k_ are the 2-dimensional mean vector and 2 × 2 covariance matrix associated with state *z_k_*. The full definition of the Hidden Markov Model is given by (*A*, {*μ_k_*, *Σ_k_*; *k* = 1, …, *K*}, *π*). These parameters were fit through iterative re-estimation using the Baum-Welch algorithm.

#### 2.6.4 Hidden Semi-Markov Models

An assumption of HMMs is that the state at time t is conditionally independent of all other states and observations given the state in the previous time step,

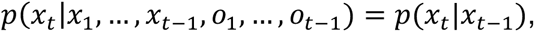

where *x_t_* and *o_t_* denote the state and observation at time *t*. However, when the hidden state of the system being modeled depends on more than just the previous time step (e.g., when the time to transition from one state to another is longer than one sample period), we can expand our hidden state model to a Hidden Semi-Markov Model (HSMM). This family of models can capture time inhomogeneity of state transitioning behavior and has been used previously in neuroscience to account for non-exponential state dwell times [44–46].

Formulations of HSMMs can include state transition rates that are functions of linear combinations of past observations and/or stimuli, unique transition probabilities for each time step, or transition probabilities that incorporate the amount of time spent in a state [45]. We utilized the *ssm* Python library [40], which implements an HSMM as an expanded Markovian state space model with limited connectivity, where multiple substates map to a single superstate:

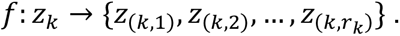

Note that *f* is a parcellating function mapping the *k^th^* superstate, *z_k_*, into *r_k_* substates. Intra-superstate transitions are only allowed in the forward direction and follow a geometric distribution,

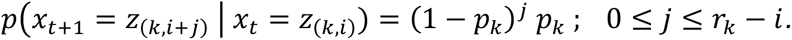

Here, *p_k_* is the probability of remaining in the current superstate. The above equation denotes the probability of moving *j* substates forward from the current, i^th^, substate between time *t* and *t* + 1. Additionally, the modeled system may move to the first substate of another superstate at any time (from the i^th^ substate of the current superstate) with probability

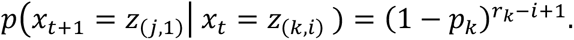

The HSMM is therefore fully defined by (*K*, {*r_k_, p_k_, μ_k_, Σ_k_* ; *k* = 1, …, *K* }, *π*), where K is the number of superstates. By including the internal dynamics of states, the model accounts for amount of time spent in a state. Geometric internal dynamics yield state durations that follow a negative binomial distribution. Parameters were fit through iterative re-estimation using the Baum-Welch algorithm.

#### 2.6.5 Hyperparameter optimization of HSMMs

Given an observation model and a prespecified number of superstates (see section 3.3), the only hyperparameter left to tune is the number of substates per superstate. To optimize this hyperparameter we swept from 1 to 30 substates using 5-fold cross-validation on the training data of each animal. Distributions across animals of cross-validation performance on sample-wise and state-wise accuracy and F1 score (see section 2.8) were compared for all substate values. The optimal number of substates was chosen as the lowest value after which there was no significant increase on the given performance measure.

### 2.7 Clustering analysis

We used 10-fold cross validation of the log likelihood complemented by silhouette score analysis to determine the number of emergent clusters in the 2D principal components space. Log likelihoods were calculated for GMM fits with 1 to 5 components, and the GMM fit maximizing the log likelihood determined the number of clusters present in each recording. We also used silhouette scores (MATLAB’s *silhouette*), a measure of the similarity of all points to their own cluster relative to the nearest neighboring cluster, as a model selection tool. For each recording, silhouette scores were averaged across all data points for each GMM fit to identify the quality of distinct clusters in the data.

### 2.8 Classification analysis

We evaluated algorithm performance post hoc by comparing their cluster assignments to the ground truth proxy cortical state labels arising from behavioral videography. Since the order of clusters provided by these unsupervised algorithms change depending on random initialization during fitting, clusters were permuted to maximize alignment with behavioral labels before calculating classification performance.

We used linear discriminant analysis (LDA) to assess the similarity between the GMM clusters and behavioral labels arising from videography, essentially providing an upper bound on the unsupervised performance when evaluated in the context of behavioral labels. To do so, we fit an LDA model on the first half of each recording and their classification performance was tested on the second half of each recording (50% training data, 50% testing data). LDA performance was quantified and compared to unsupervised two-component GMM fits by the area under the receiver operating characteristic curve (AUROC). The LDA performance on classifying behavioral labels represents an upper bound on the linear separability between whisking versus quiescent data points because it is a supervised method.

For comparing the classification performance of unsupervised algorithms to each other we first quantified accuracy and F1 scores [47] relative to behavioral labels on a sample-wise basis, where every sample was considered a prediction. However, to penalize the number of spurious state transitions, we also quantified classification performance on a state-wise basis, where each contiguous state period was considered a prediction. Periods were determined to be false positive or false negative if the estimator incorrectly predicted the behavioral label over the entire epoch, otherwise they were considered true positive or true negative. The same quantifications were used as for the sample-wise analysis (accuracy and F1 score), but now summing over all prediction periods rather than individual samples. The overall effect of considering state-wise classification performance is to penalize spurious, short time-scale misclassifications of cortical state.

### 2.9 Real-time implementation

Because the probability of being in a given state at the current time point for each of our models is dependent only on current and past activity, it is possible to use these models to infer cortical state causally, allowing for implementation in real-time. We developed a causal buffer-based method that takes advantage of a quick (<1 min) offline model fitting period before real-time inference. Models and spectral features were fit using the *ssm* and *scikit-learn* libraries of Python as described above, and the real-time algorithm was implemented with a custom-written Real-Time eXperiment Interface (RTXI) [48] program with a 2 ms real-time period. To take advantage of the models and features in their native language, we embedded Python functionality into our core C++ code using CPython. For computational efficiency we reimplemented the Viterbi algorithm in C++ using Armadillo [49] for all linear algebra.

Raw electrophysiology was passed through a fourth order Butterworth 200 Hz low-pass anti-aliasing filter before being down-sampled to 500 Hz and streamed into a 512 ms (256 sample) buffer. This buffer size was chosen for sufficiency to resolve the lowest frequencies of our signal and for efficient calculation of the FFT using FFTW3 [50]. A Hamming window was applied before transforming the signal to the frequency domain with a 256-point FFT, taken every 40 ms. FFTs of the current timepoint, along with those from previous time points, were stored for temporal context for estimating the Markov chain. Because of computation time constraints of the real-time algorithm, a finite memory for observations was implemented. New samples were pushed to the back of the observation vector up to a length of 50, at which point the earliest samples were popped off the front of the vector each time a new sample was added. The amplitude spectra of the current and previous timepoints of the signal were then Z-scored and projected onto spectral features. The resulting observations were passed into an HSMM for final state inference using the Viterbi algorithm.

#### 2.9.1 Real-time emulation experiments

To directly compare our real-time and offline results, real-time experiments were emulated by loading previously collected data into RTXI using a custom electrophysiology “playback” module. The playback module continuously streamed LFP data into the real-time environment every 2 ms, as if the data were coming from a hardware filtered and down-sampled electrophysiology rig. Using our real-time preprocessing parameters, we fit spectral features and 2D HSMM models offline to simulate the data collection and fitting period of real-time experiments. We created playback files for each recording we used in offline analysis and, utilizing the HSMM models fit for each recording, performed real-time experiments in RTXI.

When using an H[S]MM to decode a Markov chain offline, the Viterbi algorithm decodes the entire Markov chain at once by finding the most likely sequence of states that led to the data. Therefore, even though the probabilities of each state are determined causally, the Viterbi decoding algorithm is acausal. However, for the final time step in the chain, the likelihood given by the Viterbi algorithm is causal, since no future data are used. Therefore, in our real-time emulation experiments we were able to causally infer the cortical state by applying the Viterbi algorithm to current and past data only, using the latest timepoint as the final step in the Markov chain.

### 2.10 Data and code sharing

Code for the RTXI module of the real-time implementation of our method is available at: https://github.com/CLOCTools/rtxi-StateInferenceEngine. The data from this study along with analysis code are available on Zenodo: https://doi.org/10.5281/zenodo.8057803.

## 3. Results

To investigate the dynamics of cortical state switching during wakefulness, we recorded spontaneous neural activity from the whisker region of primary somatosensory cortex (S1) of 10 awake, head-fixed mice (Fig. 1a). Recordings were made using silicon multi-electrodes, and the resultant signals were bandpass filtered and down-sampled to obtain the local field potential (LFP). Laminar identification was utilized to locate putative S1 layer 4 (L4) channels (see section 2.3), and all analyses were conducted on the central L4 channel, which was found to be maximally informative of state-dependent sensory processing in previous studies [51]. Simultaneously, we recorded videography of the whiskers to identify periods of whisking and quiescence (Fig. 1b, see section 2.4). Behavioral markers of active exploration are strongly correlated with cortical state, as shown for both whisking [13,18] and locomotion [19,37], and they are also highly correlated with one another [17]. Therefore, in post hoc analyses, we evaluated the unsupervised LFP-based cortical state classifications in the context of whisking versus quiescent behavioral labels.

**Figure 1.**
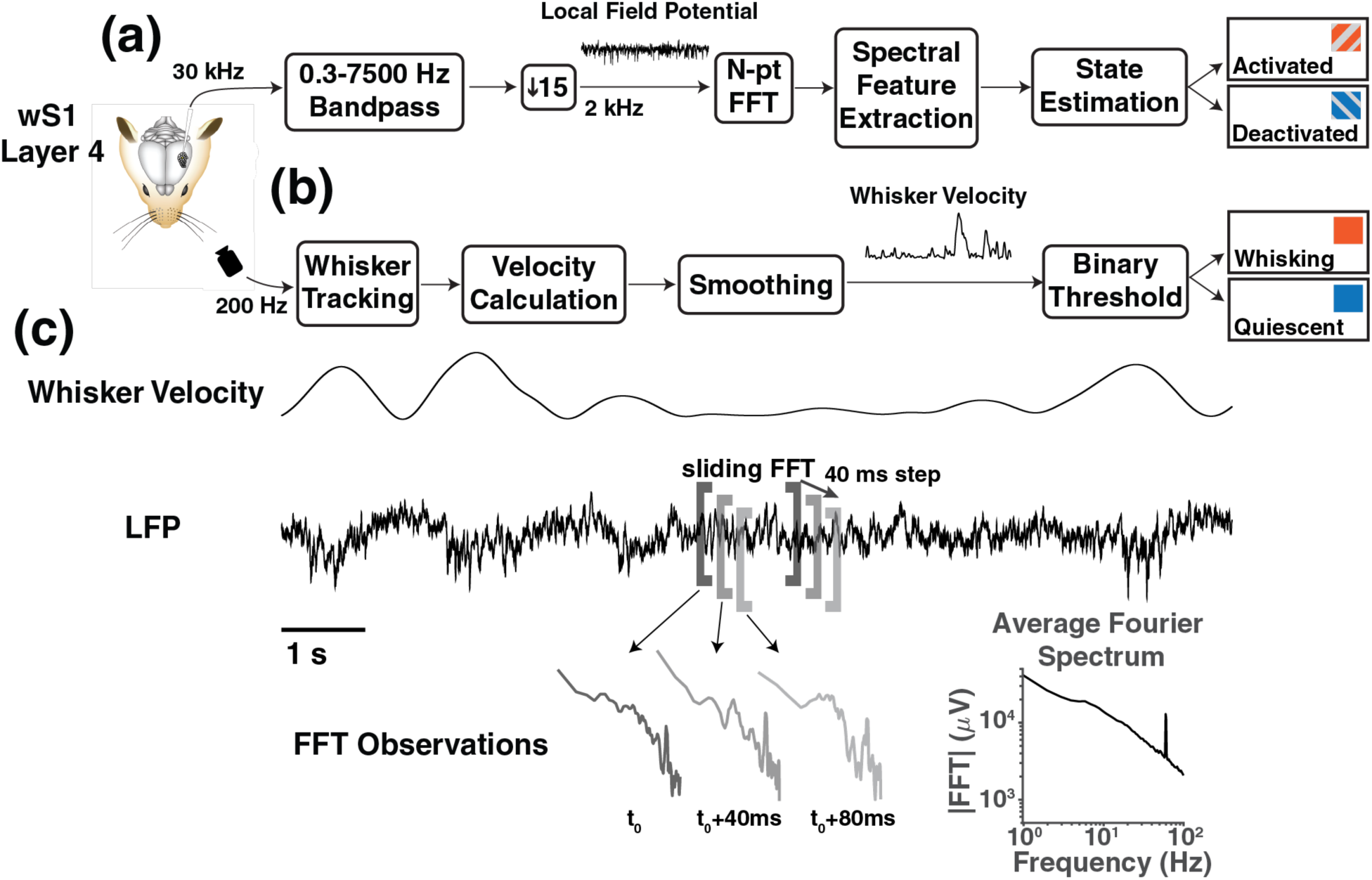
Extraction of Electrophysiological and Behavioral Signals. **(a)** Local field potentials (LFPs) were recorded from cortical layer 4 (L4) of mouse primary somatosensory cortex (S1) and processed into spectral features for cortical state estimation. **(b)** Whisker velocity traces were extracted from facial videography of the snout and a threshold was applied for behavioral state identification. **(c)** An example snippet of concurrent whisker velocity and LFP traces. A 1-s sliding window FFT was applied to the LFP signal every 40 ms, resulting in uniformly sampled Fourier amplitude spectrum observations.

As an overview of the study, we used uniformly sampled Fourier spectra of the LFP as the signal for classifying cortical state from neural activity (Fig. 1c, see section 2.3). We identified natural and emergent electrophysiological clusters, or cortical states, through unsupervised dimensionality reduction with PCA and clustering with Gaussian Mixture Models (GMMs) (Fig. 2). In post hoc analyses, we found that these emergent cortical states closely resembled clusters labeled in accordance with behavioral indicators (Fig. 3), confirming the link between cortical activation and deactivation with whisking and quiescence. Next, we used Hidden Markov Models (HMMs) and Hidden semi-Markov Models (HSMMs) to leverage temporal dynamics and decode cortical state switching with a higher accuracy (Fig. 4). Finally, we adapted these algorithms for use in a real-time context, where cortical states were decoded through machine learning approaches from streamed-in electrophysiological signals (Fig. 5).

**Figure 2.**
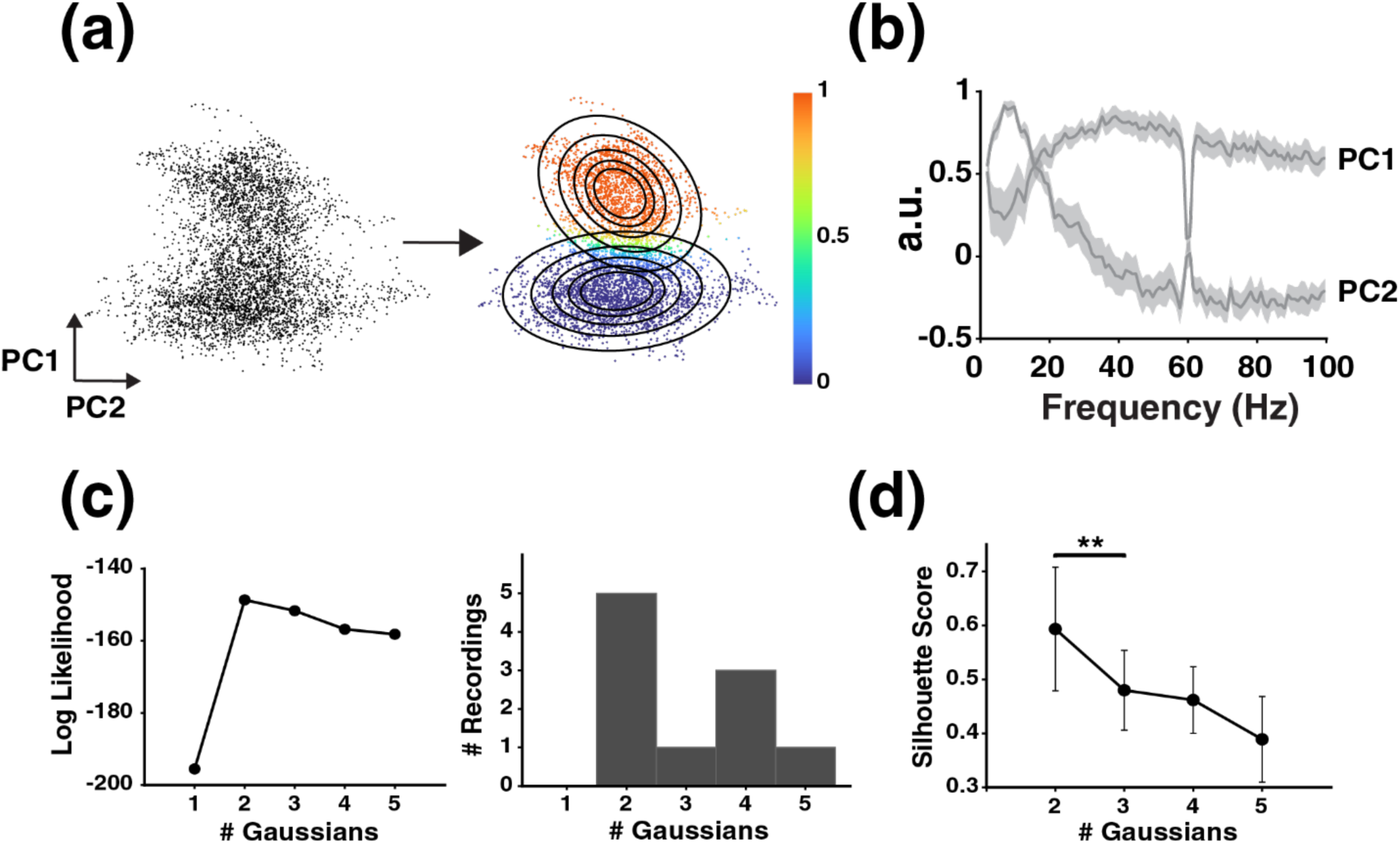
Emergent Clusters in a Low Dimensional Principal Components Space. **(a)** A depiction of a 2-component Gaussian Mixture Model (GMM) fit. Each point represents a single Fourier amplitude spectrum mapped onto a 2D principal components (PC) space. The color bar indicates the probability that a given point is assigned to the top Gaussian cluster. **(b)** The first 2 PCs (mean ± SEM) across all N=10 recordings. **(c)** (Left) The mean log likelihood of GMM fits ranging from 1 to 5 components for a single representative recording, using 10-fold cross validation. (Right) A histogram showing the number of Gaussians maximizing the cross-validated mean log likelihood across all recordings. **(d)** Silhouette scores (mean ± SD) across all recordings for cluster assignments arising from multicomponent GMMs. One-sided Wilcoxon signed-rank test; **p < 0.01.

**Figure 3.**
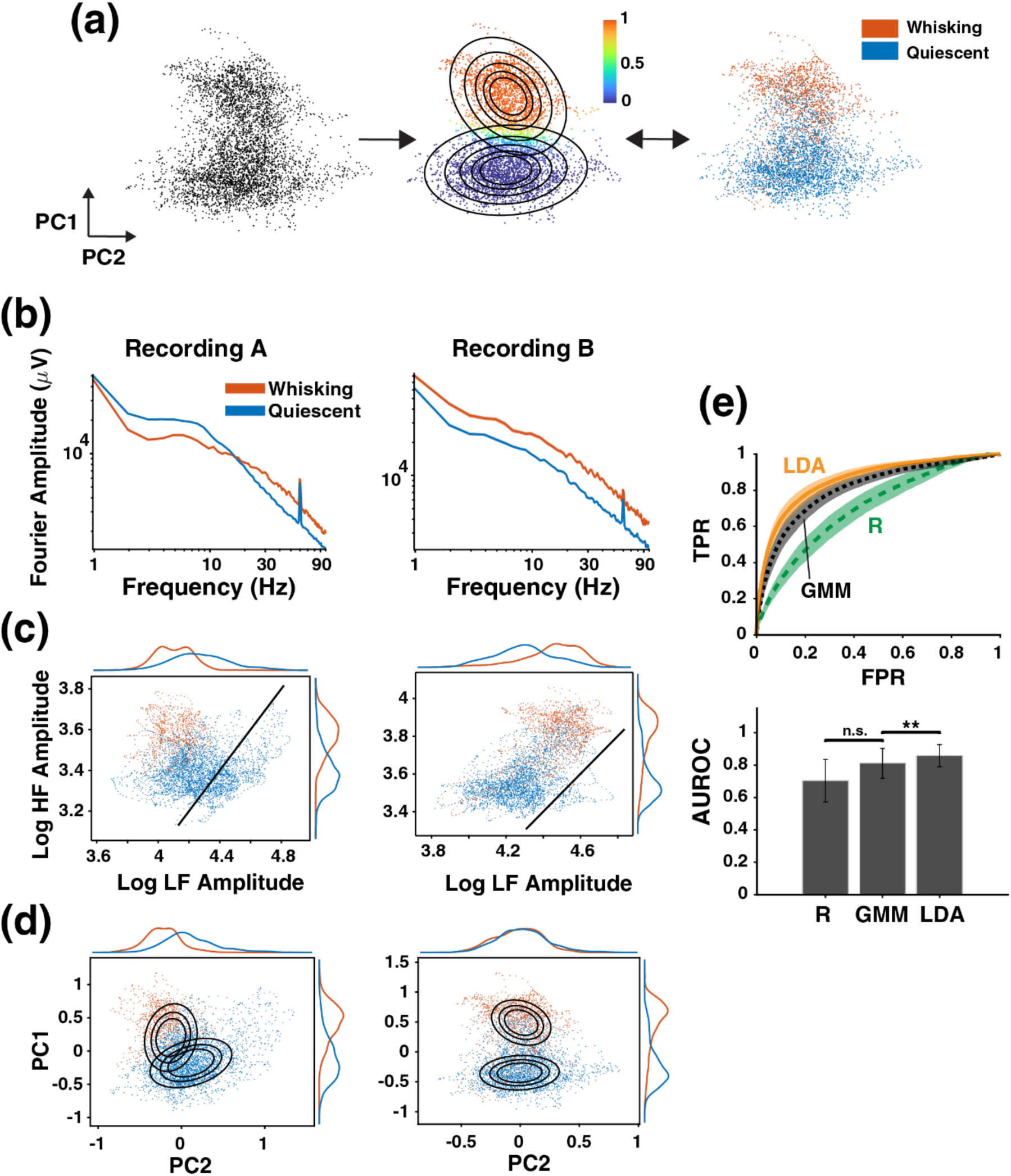
Unsupervised Gaussian Mixture Model Clusters Recapitulate Behavioral Clusters. **(a)** Comparison of a 2-component Gaussian Mixture Model (GMM) fit with behavioral labels arising from videography. The color bar indicates the probability that a given point is assigned to the top Gaussian cluster. **(b)** Fourier magnitude spectra (99% CI of the mean) separated by whisking (orange) versus quiescence (blue) for two example recordings. **(c)** Behaviorally labeled scatter plots in the 2D (log-log) LF vs. HF space for two recordings (same as in b). Marginal distributions are plotted above (LF) and to the right (HF) of each plot. The black solid line indicates the boundary for a threshold of LF/HF=10, or R=0. **(d)** Behaviorally labeled scatter plots in the 2D PC space, with overlaid Gaussians coming from a 2-component GMM fit. Marginal distributions are plotted above (PC2) and to the right (PC1) of each plot. **(e)** (Above) The averaged ROC curves (mean ± SD) for the log-transformed ratio R as well as unsupervised GMM and supervised LDA fits. GMM and LDA models were fit on data from the 2D PC space and evaluated on held-out data samples in a testing set. Below are the corresponding AUROCs (mean ± SD). One-sided Wilcoxon signed-rank test with Bonferroni correction for two comparisons; **p < 0.01.

**Figure 4.**
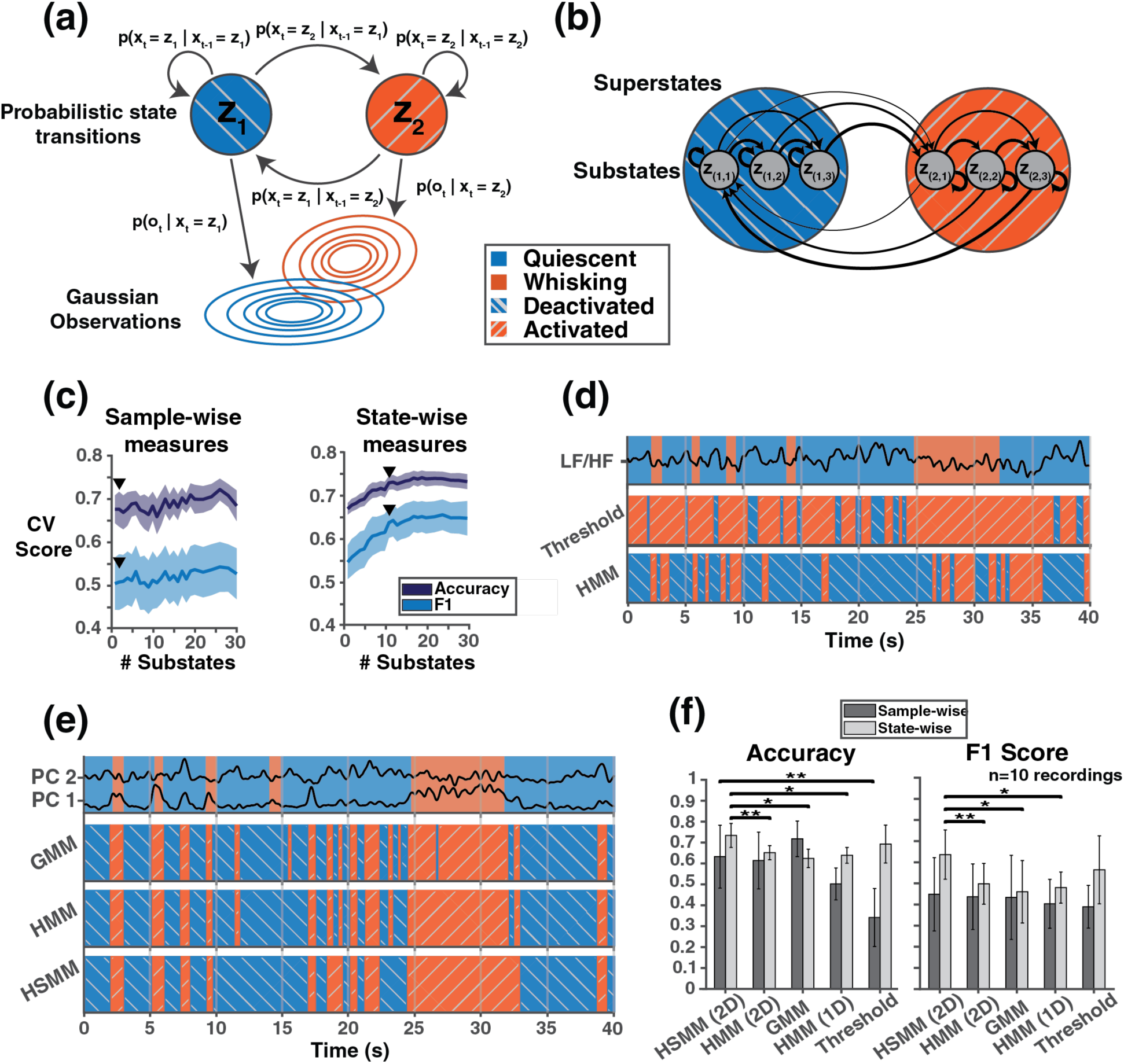
Leveraging Temporal Dynamics Increases Model Performance. **(a)** Graphical illustration of a 2-state Hidden Markov Model (HMM) with Gaussian observations. Each state has associated probabilities of staying in the same state or transitioning to the other state at the current timepoint. These transitions are conditionally independent on all previous states and observations given the current state. At each timepoint the system generates some observation, the distribution of which depends on the current state. **(b)** A Hidden Semi-Markov Model (HSMM) implemented as an expanded state space with limited connectivity by mapping the original two states, now “superstates” (blue, orange), to 3 substates (gray) each. Black arrows indicate allowed transitions and thicker line weights reflect higher probability of a transition. As the model gets deeper in a superstate the likelihood of transitioning to the other superstate increases. Transitions entering a superstate are only allowed to the first substate. **(c)** 5-fold cross-validation scores sweeping across the number of substates per superstate for the HSMM yield mostly flat sample-wise measures but significant performance gains for state-wise measures. Black arrowheads demarcate the lowest substate value at which no significant differences were found relative to all higher numbers of substates on the given measure (p > 0.05, one-sided Wilcoxon signed rank test with Bonferroni correction for multiple comparisons). Cortical state estimations (activated, orange with hash; deactivated, blue with hash) of all models are depicted during a 40 second snippet from an example recording using **(d)** 1D models acting on the LF/HF ratio and **(e)** 2D models acting on the first two principal components jointly, alongside behavioral labeling of whisking (orange) and quiescence (blue). Thresholds were fixed at LF/HF = 10 (or R = 0 in log space) for all animals. Increasing the time horizon (GMM < HMM < HSMM) reduced the number of brief misclassification periods, most often false positive. **(f)** Summary measures show the 2D HSMM outperforms all other models on at least one metric. One-sided Wilcoxon signed-rank test with Bonferroni correction; *p < 0.05, **p < 0.01.

**Figure 5.**
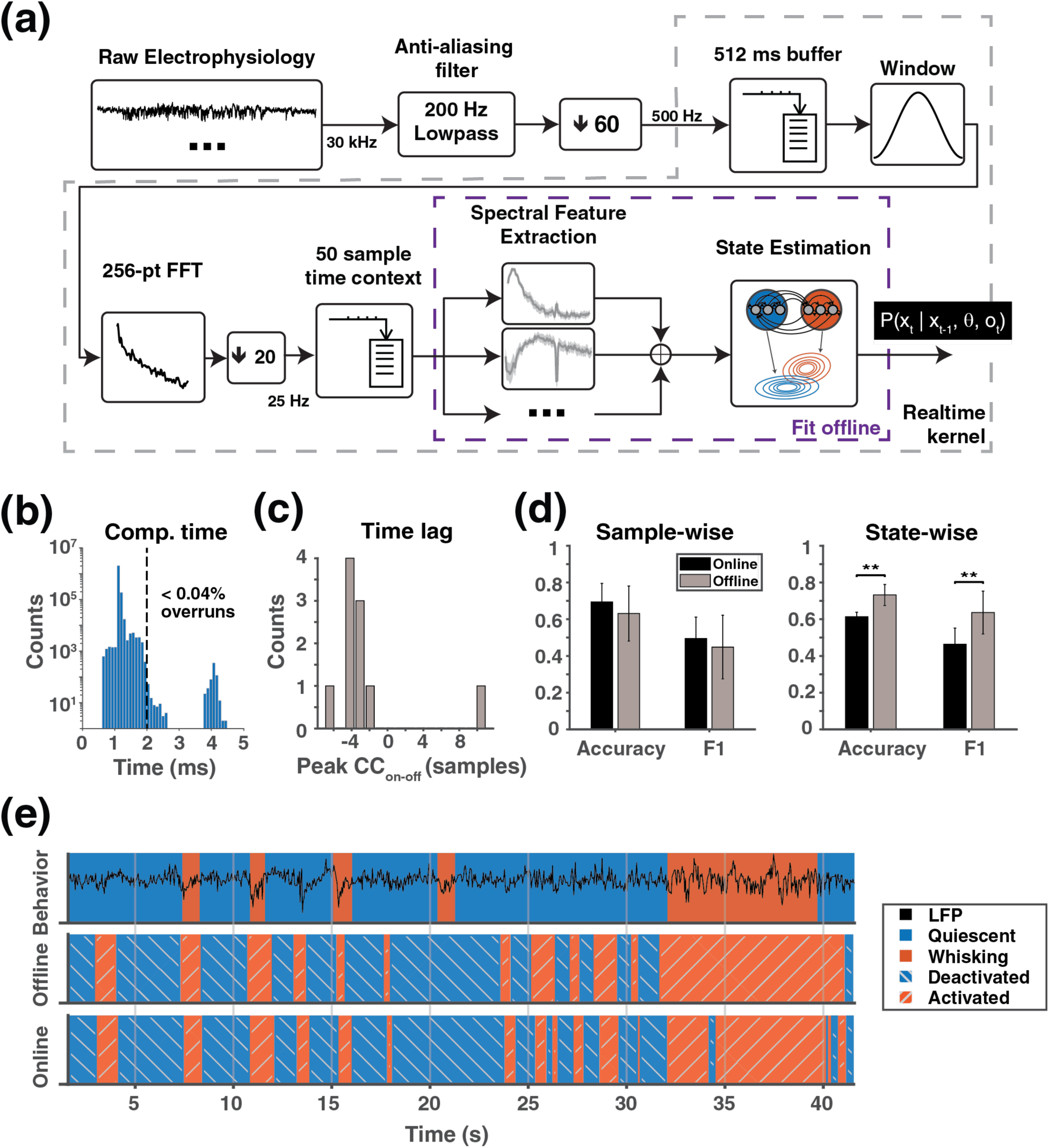
Real-time Implementation of State Estimator. **(a)** Causal buffer based real-time block diagram. Incoming electrophysiology is filtered and down-sampled to produce local field potentials (LFPs) that are loaded into a 512 ms buffer. Incoming signals to the real-time kernel are 500 Hz, which sets the real-time sampling interval at 2 ms. LFPs are transformed to the frequency domain and spectral features are extracted and used for state estimation. Spectral features and state estimator are fit offline. **(b)** Closed loop computation time across all recordings on a log scale. 99.7% of all loops finish before the ideal real-time sampling interval of 2 ms (dashed black line). **(c)** Peak cross correlations between online and offline predictions (CC_on-off_) reveal that online estimates tend to lag offline estimates by ∼3-6 samples (120-240 ms). **(d)** Summary measures of the online estimator’s performance show equivalence to the offline implementation in sample-wise metrics but moderate reduction on state-wise accuracy and F1 scores. One-sided Wilcoxon signed ranked test, **p < 0.01. **(e)** Online (bottom) and offline (middle) estimates of activated (hashed orange) and deactivated (hashed blue) cortical states next to behavioral labels (top) of whisking (orange) and quiescence (blue) for a 40 second period from an example animal with the LFP (black) overlaid on behavior.

### 3.1 Emergent clusters in a low dimensional Principal Components space

We first sought to identify emergent clusters in the spontaneous electrophysiological data which would indicate the presence of discrete cortical states. We used PCA to project each high-dimensional FFT observation into a 2D PC space, then used unsupervised GMM fits to identify emergent clusters in the data (Fig. 2a). A dimensionality reduction step was required prior to clustering to constrain the number of GMM parameters to fit. The PC features used to project into this low-dimensional space showed similar structure across recordings, with the first PC resembling a high-pass filter from 20-100 Hz and the second PC resembling a low-pass filter with a peak at 10 Hz (Fig. 2b). Interestingly, these filter characteristics are similar to the HF (30-90 Hz) and LF (1-10 Hz) bands used for traditional cortical state estimation [13]. A regression analysis confirms that the first two PCs quantitatively resemble these idealized high- and low-pass filters (Supplemental Fig. S1).

For each recording, we fit GMMs with 1 to 5 components to identify the number of discrete clusters occurring in the electrophysiological data. Shown in Figure 2a is the GMM fit for an example dataset using 2 components, where the color bar represents the probability of each datapoint belonging to the upper Gaussian. Because GMMs provide a statistical model of the data, we first used the log likelihood to evaluate the goodness-of-fit of each model to the data. Using 10-fold cross validation, we found that the log likelihood was maximized at two or more Gaussians across all recordings, with 50% of the recordings indicating a best fit with two Gaussians (Fig. 2c, right). A plot of the log likelihood versus number of Gaussians for a typical recording is shown in Figure 2c (right). The log likelihood increases substantially after a single Gaussian before reaching its maximum at some small number of Gaussians (e.g., 2), then begins to decrease for higher numbers of Gaussians due to overfitting. A silhouette score analysis confirms that the data are best explained by two clusters. The clusters arising from higher-component GMMs have significantly lower silhouette scores than two-component GMMs (Fig. 2d), indicating that additional clusters are not well isolated from the dual principal clusters in the data. Because calculating silhouette scores requires a neighboring cluster, it cannot be evaluated for a one-component GMM fit. However, the log likelihood cross validation analysis clearly indicates that a single cluster does not suitably explain the multimodal nature of the data. Taken together, these results indicate that there are two principal clusters that emerge from the spectral properties of the LFP for the data collected under these conditions, consistent with previous literature indicating discrete periods of cortical activation and deactivation during wakefulness [13].

### 3.2 Unsupervised Gaussian Mixture Model clusters recapitulate behavioral clusters

To confirm that the two discrete clusters arising from the electrophysiology correspond to cortical activation and deactivation patterns, we compared the emergent clusters arising from 2-component GMM fits with behavioral labels of whisking and quiescence. Unsupervised GMM fits did not use behavioral labels, yet we found good qualitative agreement between the GMM-based clusters and the behaviorally labeled clusters arising from videography (Fig. 3a).

In accordance with the literature, we found an observable difference in Fourier amplitude spectra when an animal was whisking as compared to when it was quiescent (Fig. 3b). In some recordings, periods of cortical activation had reduced low frequency (LF, 1-10 Hz) content and enhanced high frequency content (HF, 30-90 Hz) as has been reported previously (Fig. 3b, left) [13,19]. However, in other recordings, periods of cortical activation had increased frequency content broadly across the whole physiological frequency range from 0-100 Hz (Fig. 3b, right). The difference between these two recording types is also apparent in behaviorally labeled scatter plots in the log HF versus log LF space, by the relative orientation of the whisking cluster compared to the quiescent cluster (Fig. 3c). Reduced LF content and enhanced HF content during whisking produces clusters that are perpendicular to the identity line in this space (Fig. 3c, left), whereas a broadband increase in frequency content during whisking produces clusters that are oriented along the identity line (Fig. 3c, right). These distinct recording “phenotypes” likely reflect underlying physiological variability in the presentation of cortical states, underscoring the need for cortical state decoding tools that are robust across different animals and recording sessions.

Interestingly, we can plot the decision boundary of the log-transformed LF/HF ratio (R = 0) as suggested by the literature [13,18] in the log HF versus log LF space, which does not separate the behaviorally labeled clusters well. Even when sweeping the specific threshold value on R—which would correspond to translating the decision boundary through the log HF versus log LF space—it only provides optimal discrimination in the case of a perfect anticorrelation between LF and HF. On the other hand, unsupervised dimensionality reduction into the 2D PC space and unsupervised GMM fits provided much better alignment with behavioral clusters across recording phenotypes (Fig. 3d). This is quantified in Figure 3e across all recordings, where we report AUROCs ranging from 0.66-0.93 for GMM fits and 0.40-0.82 for thresholds on R. This does not rise to the level of significance (Fig. 3e, bottom; p=0.08, one-sided Wilcoxon signed-rank test) because R better separates the behavioral clusters in 2 out of 10 recordings. We additionally report that the unsupervised GMM fits perform nearly as well as an optimal, linear classification boundary in the 2D PC space arising from linear discriminant analysis, which had AUROCs ranging from 0.71-0.94 (Fig. 3e, bottom). This supervised approach operates on behavioral labels that do not inform the GMM fits, and therefore represents an upper-bound of performance for linear methods in a given projection space. Altogether, these results suggest that the emergent clusters arising from the electrophysiology recapitulate behaviorally labeled clusters. As such, we proceed by denoting the two emergent clusters from unsupervised approaches as activated and deactivated cortical states, due to the noted high correlation between these cortical states and whisking.

Importantly, we found that behavioral clusters were highly separable in the 2D PC space, indicating that it was an appropriate space to find inherent cluster structure in the data. In particular, behavioral clusters were more separable in the 2D PC space compared to using the full Fourier amplitude spectra and comparably separable to the 2D log HF versus log LF space (Supplemental Fig. S2a). Moreover, behavioral clusters were not separable for PCs after the second component, indicating that keeping the first two dimensions was sufficient for separating out cortical states (Supplemental Fig. S2b-c).

### 3.3 Temporal dynamics of state transitions with Hidden [Semi-]Markov Models

The methods outlined above, thresholding and GMMs, do not take advantage of temporal information; the likelihood of being in either state is determined by the current observation only. GMMs simply find the maximum *a posteriori* cluster labels given a Gaussian prior and some number of clusters. In effect they find the optimal decision boundary between clusters in the feature space assuming the data are drawn from multiple Gaussians. However, the electrophysiology data modeled here is continuously streamed, sequential, noisy data (Supplemental Movie 1). We therefore incorporated state transitioning as a stochastic process into our models with a Hidden Markov Model (HMM) (Fig. 4a), which is well suited for this type of data. HMMs explicitly model an underlying state that gives rise to different distributions of observations. In this formulation, the underlying state is directly analogous to cortical state and observations are either the Gaussian distributed 2D transformations of the Fourier spectra, as in the previous section, or the 1D log transformed LF/HF ratio, R. The HMM structure captures probabilities, given that the cortex is in a particular state, of remaining in that state or transitioning to another state, as represented in the illustration in Figure 4a.

We found that behavioral state duration distributions did not follow exponential curves (Supplemental Fig. S3), but exponential state durations are characteristic of Markov processes that are inherent to the HMM framework [41]. This finding indicated that cortical state transitioning is not truly a Markov process and therefore the transition dynamics incorporate more than just the previous timepoint. Since all the information needed to determine the likelihood of the system being in a particular state is not summarized in a single timepoint, it seems that transitions occur over a time period that was longer than a single sampling period, which was equivalent to the FFT window step length (40 ms, see section 2.3). Therefore, we explored whether a Hidden Semi-Markov Model (HSMM) (Fig. 4b), a common extension to HMMs for accounting for non-exponential state dwell times [40,44–46], could better explain the data (see section 2.6.4). Consistent with our finding in the previous section that a mixture of two Gaussians best fit the data, we utilized 2 state HMMs and 2 superstate HSMMs with Gaussian observation models for each animal.

Our HSMM was implemented as an expanded state space with limited connectivity [40], where multiple substates map to each superstate (Fig. 4b). In this formulation, superstates are analogous to states in an HMM and substates are intermediate steps in the state switching process. Substates and their associated transition probabilities can therefore be thought of as modeling the internal dynamics of state switching. Intra-superstate transitions can only go in the forward direction and inter-superstate transitions can only go to the first substate of the given superstate as depicted in Figure 4b (see section 2.6.4). Given the above observation model, the only free hyperparameter to tune is the number of substates per superstate. Using 5-fold cross validation on HSMMs with 2D observation models, we swept the number of substates from 1 to 30. Because models with higher numbers of substates can at worst reproduce the behavior of those with lower numbers of substates, we found that adding more substates could only improve the models’ performance up to a saturation level. We therefore determined optimality by finding the lowest number of substates that was indistinguishable from more complex models on all performance measures using a one-sided Wilcoxon signed rank test with a Bonferroni correction for multiple comparisons. We found that sample-wise accuracy and F1 score reached equivalence by 2 substates while state-wise accuracy and F1 score did not reach equivalence until 11 substates (Fig. 4c, see section 2.8 for description of metrics). Therefore, we took the optimal number of substates per superstate to be 11, since it was the lowest value to achieve maximum performance on all metrics. An optimal substate value of 11 per superstate taken together with a sampling period of 40 ms indicates that the time period that best captures the dynamics of cortical state transitions is ∼440 ms.

With optimal parameters of each model chosen, we sought to characterize estimates from each model. Both 1D estimators (binary thresholding at R = 0 and 1D HMMs with R observations) were biased toward activated cortical states, as exemplified in Figure 4d. Indeed, for all animals the 1D models provided biased estimates (Supplemental Fig. S4a-b). However, the threshold of R = 0 was taken from the literature [13,52] and not fit for each animal. Even so, using a different threshold is unlikely to eliminate this effect, as the 1D HMMs, which were fit to each animal, were still skewed toward activated cortical states. This suggests that bias arises because of the simplicity of the LF/HF ratio, which is supported by a marked decrease in bias of the 2D models (Supplemental Fig. S4a-b). However, the 2D models were susceptible to noise while close to the boundary between clusters, as evidenced by brief misclassifications in Figure 4d. Generally, their most common errors were brief false positives, or periods during quiescent behavior predicted as activated cortical states (Supplemental Fig. S4b). Of the 2D models, the GMMs most often erroneously predicted brief periods of activation during quiescence. These errors were reduced as the retrospective time horizon of the models increased (from 0 ms) to 40 ms in the HMMs and 440 ms in the HSMMs. An increasing time horizon had the effect of smoothing the models’ state estimation and making them more robust to measurement noise, especially close to the decision boundary. One notable trend of the 2D models is that predictions of activated cortical states seem to precede whisking behavior. It is unclear whether cortical states truly precede behavioral states or if this is a symptom of model bias.

Estimator bias and general agreement with behavior were most prominently reflected in the sample-wise measures while resilience to noise was reflected in the state-wise measures. Across all animals, HSMMs with 2D observation models attained the best overall performance across sample-wise and state-wise accuracy and F1 score (see section 2.8), achieving significantly better scores than all other models tested on at least one metric and never worse than any other model (Fig. 4d-f). Because state-wise accuracy and state- and sample-wise F1 scores renormalize based on the amount of predicted positive class (here we considered *activated* and *whisking* as positive), sample-wise accuracy is most appropriate for assessing large disagreement in labels created by bias. This is evident in the dramatically worse sample-wise accuracy of the thresholding method compared to the 2D methods. For more subtle differences in the models, it is instructive to compare state-wise accuracies, in which brief, erroneous state switches are reflected, and state-wise F1 scores, which provide a measure of set similarities. A high F1 score indicates that the sets are both overlapping and of similar size. On these state-wise metrics the 2D HSMM outperforms the other 2D models and the 1D HMM, indicating better robustness to noise and higher similarity to the behavioral labels. This robustness without the need to temporally smooth measurements or average across multiple trials makes the 2D HSMM the most viable unsupervised algorithm for cortical state decoding.

### 3.4 Real-time cortical state estimation

Taking advantage of the causality of H[S]MMs, we implemented a causal buffer based real-time state estimator (Fig. 5a) to infer the state at the current timepoint. This system takes in a continuous stream of electrophysiological data, passes it through an anti-aliasing filter and down-samples to obtain LFPs, and writes the data into a 512 ms buffer. Inferences are obtained as in the offline case through conversion to the frequency domain, projection onto spectral features, and decoding the sequence of states through the Viterbi algorithm, with the notable caveat that the features and model were fit offline. Additionally, because of computational constraints imposed by real-time requirements, a finite memory of 50 FFT samples was implemented (i.e., finite memory of 50 × 40ms = 2 seconds). This finite memory limited the models’ context for decoding the most likely sequence of states. For direct comparison of the offline and real-time implementations, we re-analyzed our previously recorded data through real-time emulation experiments, where we streamed data as if it were being read in from an ADC board. It should be noted that while state transition probabilities are defined causally in H[S]MMs, the Viterbi algorithm [43] decodes the entire sequence of states in all available timesteps at once, and so in general is not causal. However, if only past and present data are fed into the algorithm the state inference of the online implementation at the current timepoint is causal, in contrast to the offline implementation.

With a sampling frequency of 500 Hz, the real-time sampling interval of the system was 2 ms. Across all animals the real-time implementation had mean latencies ranging from 1128.1-1168.0 μs with variances between 393.0-928.3 μs and closed loop latencies were less than 2 ms in 99.97% of iterations (Fig. 5b). Larger than expected variances were a result of a bimodal latency distribution with a small fraction of iterations (< 0.04%) taking much longer than others. Compared to behavioral labels, sample-wise accuracies and F1 scores across animals were 0.69 ± 0.10 (mean ± std) and 0.50 ± 0.12, while state-wise accuracies and F1 scores were 0.62 ± 0.04 and 0.49 ± 0.10, respectively (Fig. 5d-e). For equivalent comparison to our offline method, the same model and preprocessing—most notably down-sampling to 500 Hz and 512 ms spectrogram window as opposed to 2 kHz down-sampling and 1 s spectrogram window—were applied to the data offline using the acausal Viterbi algorithm. The real-time method performed equivalently well as the offline algorithm on sample-wise accuracy (0.63 ± 0.15, p=0.19; two-sided Wilcoxon signed rank test) and sample-wise F1 score (0.45 ± 0.17, p=0.38), but performed worse on state-wise accuracy (0.73 ± 0.06, p=0.002) and state-wise F1 score (0.63 ± 0.12, p=0.002). This small drop in performance can be attributed to the lack of future context in our real-time system that is available offline.

Additionally, we sought to characterize any temporal differences between the real-time and offline estimators. Therefore, we calculated the cross-correlation between real-time and offline estimates (Fig. 5c-d). The cross-correlation analysis revealed that correlations were highest with delays of 3-6 samples for most animals, meaning that the real-time estimator lags the offline estimate by ∼120-240 ms. This result can again be attributed to the lack of future context of the real-time estimator. Interestingly, this would suggest that the offline estimator incorporated information about where the system was going to determine the time of state switch during an ambiguous period. Due to the lack of ground truth cortical state labels, it is difficult to determine which estimator more accurately timed state switches. Future work may address this by investigating the processing consequences of cortical state, such as changes of sensory evoked responses, during these uncertain regimes. However, the good estimation accuracy relative to behavioral proxy labels taken together with the fast computation time demonstrate that real-time cortical state estimation is achievable with H[S]MMs.

## 4. Discussion

Our study aimed to develop a data-driven approach for robust, real-time classification of cortical states from electrophysiological data. We found that PCA and GMMs identified *emergent* clusters from spectral LFP data, which corresponded to cortical states as reflected in simultaneously recorded whisker activity. Subsequently, we adapted our clustering approach to capture the temporal dynamics of cortical state switching with HSMMs, leading to further improvements relative to behavioral indicators of cortical state. Finally, we implemented our unsupervised cortical state estimation approach in real-time, demonstrating the feasibility of online cortical state estimation given real-time accessible LFP measurements. A central goal of ours was to provide a continuous readout of cortical state to enable causally testing state dependent sensory processing, perception, and behavior in future studies.

Although in the present study we validated our algorithms for estimating cortical state with behavioral markers, a clear distinction must be made between cortical and behavioral states. While there has been a strong correlation reported in the literature between behavior and arousal-related cortical states [12,13,17–19,23,28], behavioral state is the motor output of an animal at a certain time, and cortical state is a quality of ongoing neural activity that leads to changes in network processing [14,15,26,53]. Activated cortical states in primary sensory cortices have been shown to increase sensory coding [54] and differentially activate projections to higher cortical areas [55]. Oscillatory states in particular have been proposed to gate information flow across local circuits [56–58]. Since our method acts only on electrophysiological data and is agnostic to behavioral labels during training and inference, our state estimates should be viewed as inferences of cortical state, which should be highly correlated with behavioral states but not necessarily one-to-one. However, ascending cholinergic activity from the basal forebrain causes cortical activation [37] and has been associated with locomotion, which is highly coincident with whisking [17], through the mesencephalic locomotor region (MLR) [59]. Stimulation of the MLR can reliably induce activated cortical states [60] and locomotion [61–63]. Therefore, these behavioral states and cortical states share common inputs and are strongly associated, affirming our choice to validate our models against behavioral labels.

In order to further our goal of providing continuous readouts of cortical state, we directed our development towards signals and algorithms that allow for real-time execution. There is evidence that cortical state switches are detectable from the activity of individual cortical neurons [13] or the combined population activity in cortex [52], but neural spikes are more difficult to access in real-time than the LFP. While the LFP requires only minimal hardware filtering and down-sampling, a major algorithmic bottleneck in accessing real-time, granular spiking data is spike sorting. Although there has been significant recent development of spike sorting algorithms, most of them are designed and intended for post hoc analysis of large, high channel count datasets [64]. There are some notable exceptions, including an FPGA-based approach for performing single-channel spike sorting in real-time that performs spike template matching in hardware [65]. However, the LFP constituted the ideal signal for estimating cortical states because of its known connections to states of arousal [13] and its high degree of accessibility in a real-time context. Note that it is likely the case that spiking data provide complementary information to that of the LFP [52] thus suggesting a framework in which both signals would be beneficial, but due to the practical constraints here, this was not taken into consideration. In addition to operating on a real-time signal, the algorithms underlying our cortical state inferences were fast and causal, as needed for online applications. We modeled the temporal dynamics of cortical state switching with HSMMs, where the state decoding could take place with a causal Viterbi algorithm considering only the likelihoods of present and past states. Our HSMMs imposed an appropriate smoothness on cortical state estimates given the dwell times in the identified cortical states. Importantly, we achieved this smoothing function in a real-time context, without the need to smooth measurements with an acausal window as is typical of post hoc state decoding methods [30]. In our recordings, we found that 11 substates (with a time step of 40 ms), or a temporal window of approximately 440 ms, best matched the time course of cortical state switching. When we implemented these algorithms in RTXI and tested them in real-time emulation, our round-trip computation time for cortical state inference was around 1100 μs, allowing us to decode cortical state given an incoming LFP signal sampled at 500 Hz. In addition to being feasible in real-time, our methods were all unsupervised, and therefore offer clear advantages by minimizing *a priori* assumptions about the characteristics of our extracted cortical states. Given a hypothesis of discrete or strongly multimodal brain states, our cross-validated cluster fits provided a principled way to uncover the number of cortical states, which can be a point of contention in the literature. For example, sleep-wake studies reported between 3 and 7 distinct states as informed by both electrophysiological and behavioral readouts [30,66,67]. We found that two cortical states best explained the data in most of our recordings, consistent with the two main cortical states of wakefulness that have been reported in the awake rodent model [13]. However, there is much speculation that the number and nature of these states of wakefulness will vary with a wider behavioral context [12]. Our unsupervised clustering methods provide one approach to identify these more granular cortical states of wakefulness when they emerge due to novel, complex, and/or ethological behaviors that go beyond the head-fixed preparation.

This same versatility provided by a minimization of *a priori* assumptions improves our method’s robustness to variability. We reported two recording phenotypes in our head-fixed animals, but our state decoding approaches were able to identify emergent cortical states robustly in both cases. In a subset of recordings, we found large, low-frequency fluctuations during quiescence as well as suppressed low-frequency fluctuations and enhanced high-frequency fluctuations during behavioral activity consistent with previous work [16,17,19,24]. However, we also showed a distinct recording phenotype where there was a broadband (<100 Hz) increase in spectral LFP content during active behaviors, which is not widely reported in the literature. The source of variability leading to the two recording phenotypes that we report is not clear. We cannot completely discount the possibility of subtle movement artifacts or other sources of electrical noise affecting the signal. However, it is likely that electrophysiological signatures of cortical state may present differently across animals due to inherent interindividual variability. Previous studies decoding sleep-wake brain states in an unsupervised fashion reported subtle but important inter-animal differences in the spectral content of awake brain states [30]. Despite this source of electrophysiological variability, we reported consistent separation between cortical states using our unsupervised methods, suggesting a robustness to unknown sources of variability across animals.

Furthermore, our data-driven methods are designed to be flexible enough to generalize across distinct brain areas with different signatures of brain state. Our approach of using PCA to find low-dimensional feature spaces bypassed the use of canonical LF and HF frequency bands that separate cortical states well in primary somatosensory cortex [13,18], but may have degraded performance in other cortical areas and limited utility for subcortical brain states. Global cortical activation due to movement and arousal presents differently across cortical regions, with profound suppression of low-frequency activity in somatosensory and motor areas, moderate suppression of low-frequency activity in auditory and visual areas, and a shift in peak low-frequency activity in selected frontal areas [18]. Moreover, anatomical studies show region-specific cholinergic projections from the basal forebrain, providing a mechanism for cortical states to be controlled differentially and locally [68]. While cortical brain states are principally associated with arousal and attention, brain state readouts from subcortical structures can have distinct behavioral relevance and electrophysiological characteristics. For example, beta oscillations (13-30 Hz) in the subthalamic nucleus (STN) are associated with Parkinsonian movement deficits in humans [7] and hippocampal theta-gamma coupling states have been implicated in memory formation [58,69]. For the latter, relative phase must also be taken into consideration. While phase was not considered in the current study, incorporating this information outlines an interesting avenue for future work towards a more general framework [51]. Altogether, the diversity of global and local brain states suggests the need for data-driven and unsupervised approaches to robustly identify brain states across regions.

Recognizing the diversity of brain states highlights the shortcomings of our method. Firstly, the PC features we used were identified based on their contribution to the overall variance of the data. It is possible that some brain states are identifiable only by subtler features with small contributions to the variance. These states would be overlooked by the current method. However, it may be possible to identify features with other dimensionality reduction techniques based on covariance [70] or local neighborhood structure [71] to better separate subtler states. Additionally, future feature identification should incorporate phase information, which is crucial for brain states identifiable through cross frequency phase-amplitude coupling [51,58,72–74]. Phase-amplitude analysis relies on precise calculation of instantaneous phases, which is typically executed with wavelet [58], Hilbert [75], or Gabor expansions [76] over a predefined period of interest, which would need to be adapted to the real-time streaming data context. Interesting work has been done for calculating cross frequency coupling with higher moments of the power spectrum [74] and with filter banks applied to the FFT [77] that show promise for real-time applications. Once extracted, these new features could in principle be paired with an H[S]MM for state estimation provided a good observation model can be formulated.

Overall, our work provides an updated approach to cortical state decoding with two main focuses. The first focus was using unsupervised dimensionality reduction and clustering techniques, which promise increased robustness across animals and brain areas. Moreover, such unsupervised techniques provide a principled manner for identifying novel brain states—outside of cortical activation and deactivation—which may emerge from recordings in richer behavioral contexts than head-fixation. The second focus was moving the cortical state estimation problem to the real-time context. This, too, promises to facilitate the study of brain states more broadly. For example, the real-time cortical state decoding scheme we proposed could be used in a reactive experiment where sensory stimuli are delivered in a state-dependent manner, which could be useful for studying state-based changes in sensory encoding, perception, and behavior. A similar paradigm could be used to deliver electrical stimulation when pathophysiological brain states are detected, as in deep brain stimulation (DBS) therapies for Parkinson’s disease. Finally, real-time estimates of cortical state could be used in a closed-loop context to control brain state in a reactive manner. Already, there are a handful of intracortical, thalamocortical, and neuromodulatory circuit mechanisms implicated for regulating cortical state [24–26,78,79], which could be used to actively and dynamically shape cortical state. Such a controller could be used to causally test the functional consequences of cortical states in sensory encoding, perception, and behavior.

## 5. Conclusion

This study offers a novel, data-driven approach to brain state decoding that is suitable for real-time deployment. By utilizing unsupervised dimensionality reduction and clustering models, we can key in on spontaneous, multimodal brain activity in a variety of contexts. In the present study, this allowed us to robustly decode cortical activation and deactivation across different animals and recording sessions, but in general these approaches can be extended to identify brain states in other areas with different characteristics from somatosensory cortex. Moreover, by advancing state estimation into the real-time domain, we open up new avenues for the study of brain states. Our proposed real-time cortical decoding scheme allows for state-dependent delivery of sensory stimuli, offering insights into state-based changes in sensory encoding, perception, and behavior. This framework also holds promise for applications such as closed-loop control of brain state via optogenetic or electrical actuation of endogenous neural circuitry.

## Notes

Funding: This work was supported by the National Institutes of Health (NIH BRAIN Grants R01NS104928, R01EB029857, RF1NS128896 to GBS, NIH R21NS112783 to GBS and AP, NIH R01NS115327 to AMFB, NIH T32EB025816 to DAW and AMFB); Swiss National Science Foundation postdoctoral fellowships P2ELP3_168506 and P300PA_177861 (AP). Additional support was provided to AS through startup funds from the University of Minnesota Medical School.

Conflict of Interest: The authors declare no competing interests

### Competing Interest Statement

The authors have declared no competing interest.

