## Supplemental Figures for "A Machine Learning Approach for Real-time Cortical State Estimation"

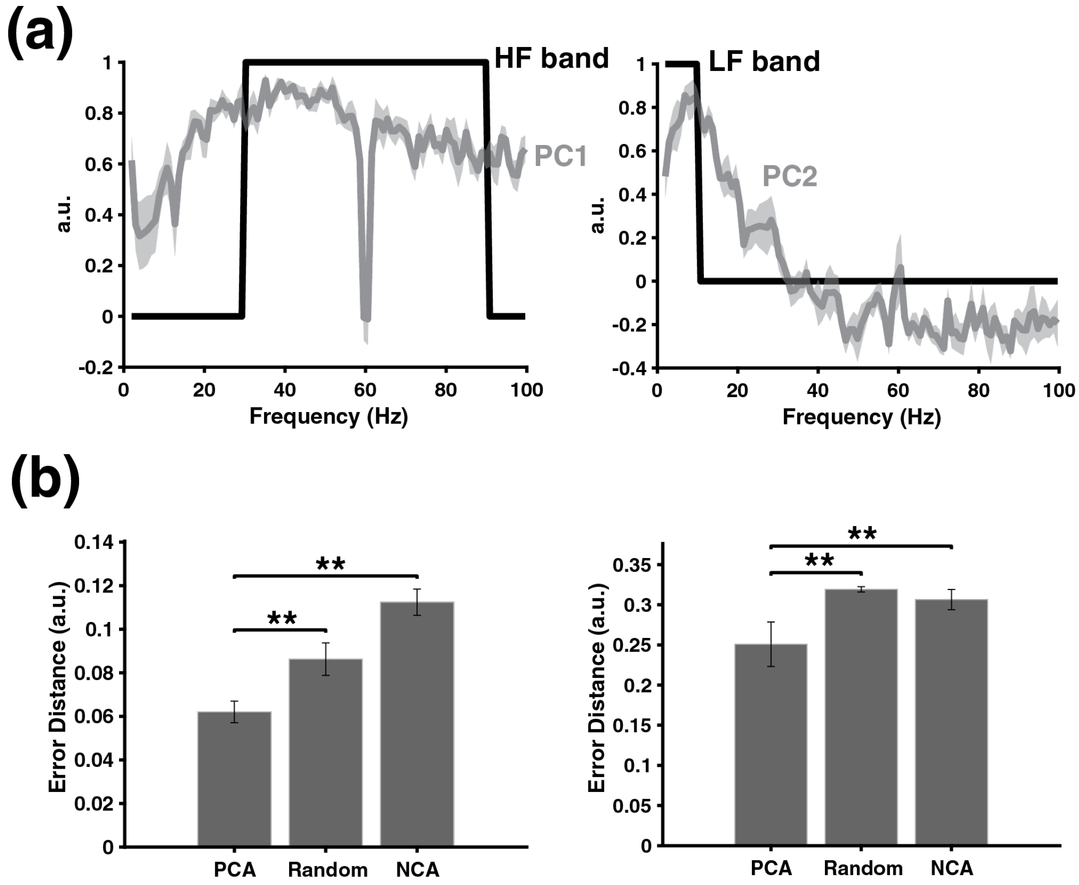


**Supplemental Figure 1)** Principal Component Features Resemble Canonical Low- and High-Frequency Bands. **(a)** (Left) The first PC (mean ± SEM) across all N=10 recordings compared to the canonical high-frequency band (30-90 Hz) indicated in black. (Right) The second PC compared to the canonical low-frequency band (1-10 Hz) indicated in black. **(b)** The error distance (mean ± SD) of linear regressions onto an idealized LF band feature (left) and HF band feature (right). From left to right, the regressors were 2D PC features, random features, and neighborhood components (NC) features. One-sided Wilcoxon signed-rank test with Bonferroni correction for two comparisons; $^{**}p < 0.01$.

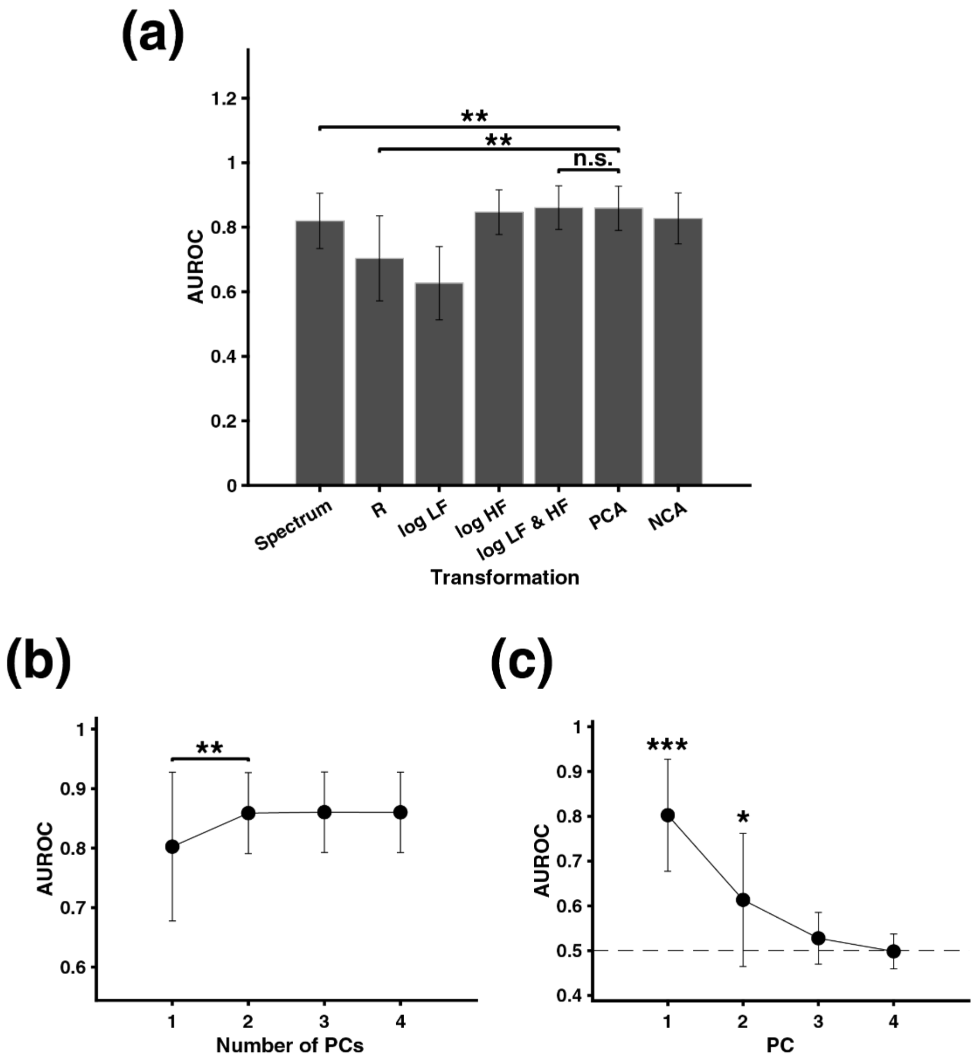


**Supplemental Figure 2)** Two Principal Components Separate Whisking and Quiescence. **(a)** AUROCs (mean ± SD) for LDA classifiers trained on different transformations of the Fourier amplitude spectra. From left to right, these transformations are the full Fourier amplitude spectra (<100 Hz, no DC), the log-transformed LF/HF ratio (R), the log-transformed LF component only, the log transformed HF component only, the 2D combination of the log-transformed LF and HF components, the 2D PCA mapping, and the 2D NCA mapping. Two-sided Wilcoxon signed-rank test with Bonferroni correction for three comparisons; $^{**}p < 0.01$. **(b)** Supervised LDA classifiers were trained on Fourier amplitude spectra data mapped to multidimensional PC spaces. Reported are the corresponding AUROCs (mean ± SD) for N=10 recordings, as evaluated on held-out data samples in a testing set. One-sided Wilcoxon signed-rank test; $^{**}p < 0.01$. **(c)** Same as in (a), except that LDA classifiers were trained on projections onto the $K^{th}$ PC only. The black, dashed line indicates chance levels of classification. One-sided Wilcoxon signed-rank test; $^{***}p < 0.001$, $^{*}p < 0.05$.

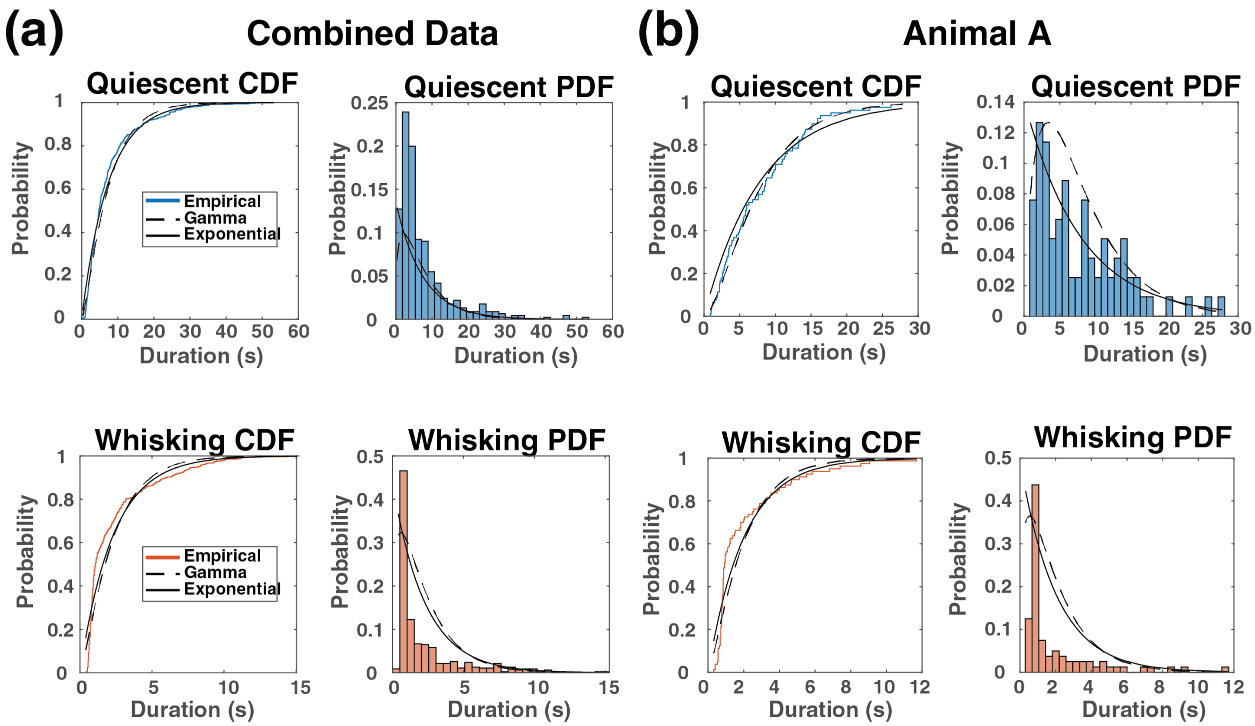


**Supplemental Figure 3)** Behavioral duration distributions are non-exponential. **(a)** Durations of quiescent (blue, top) and whisking (orange, bottom) behavioral states were aggregated across all animals. Cumulative distribution functions (CDF, left) and probability density functions (PDF, right) are depicted for each behavioral state alongside the best fit exponential (black, solid) and gamma (black, dashed) curves. Distributions of each behavioral state were found to be non-exponential ($p < 0.001$; Anderson-Darling test). **(b)** Same distributions as in (a) are depicted for a single animal. Duration distributions were found to be non-exponential ($p < 0.001$; Anderson-Darling test).

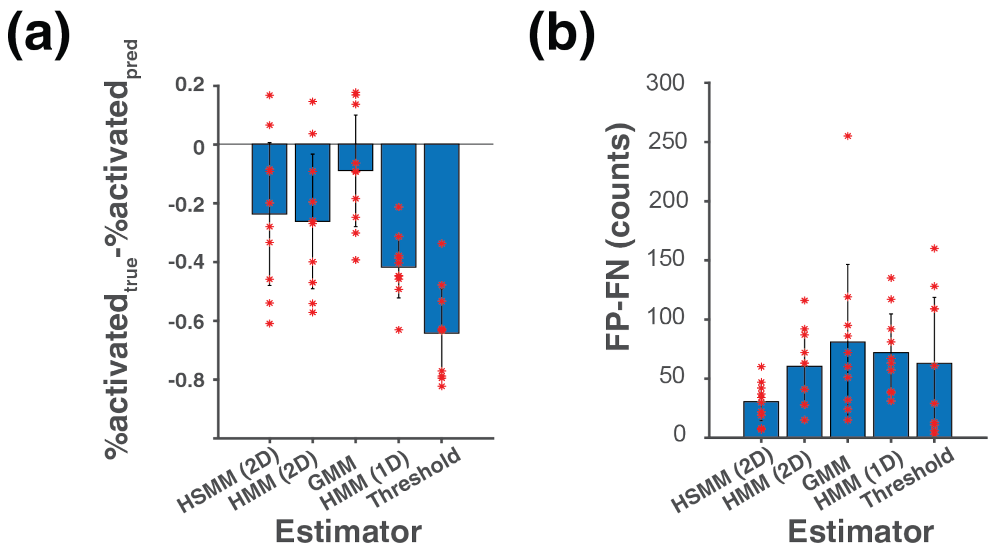


**Supplemental Figure 4)** Estimators are biased toward activated cortical states. **(a)** Bias was evaluated as the difference between the percentage of samples during which there was whisking behavior and the percentage of samples that were predicted as activated cortical states for each animal. Larger negative values of this metric indicate stronger bias toward activated cortical states. **(b)** Bias can also be calculated in a state-wise manner as the difference between false positive states and false negative states. High positive values indicate a large number of erroneous false positive periods.

**Supplemental Movie 1)** Frequency domain observations are noisy and sequential in time. **(a)** Time evolution of the Fourier amplitude spectrum (<100 Hz, logarithmic scale) for a 5 second segment of a recording is shown. Frames display the frequency domain signal in each time point, separated by 40 ms. Spectra at each time point are behaviorally labelled; they are colored orange when the animal was whisking and blue when the animal was quiescent. Considerable noise in the amplitude spectra is caused by the time-frequency uncertainty principle. **(b)** The spectra for the same timepoints as in (a) are summarized with average LF and HF amplitude (log scale). Observations are sequential in time and follow trajectories in LF vs. HF space.
